# Is bigger always better? Neither body size nor aggressive behavior are good predictors to measure the degree of specialization of hummingbird interaction networks in rocky outcrops

**DOI:** 10.1101/2021.02.27.433160

**Authors:** Ricardo Marcelino Claudino, Yasmine Antonini, Cristiane Martins, Marina do Vale Beirão, Érika Martins Braga, Cristiano Schetini de Azevedo

**Affiliations:** Programa de Pós-Graduação em Ecologia, Conservação e Manejo de Vida Silvestre, Universidade Federal de Minas Gerais, Av. Presidente Antônio Carlos, 6627, Pampulha, CEP: 31270-901, Belo Horizonte, MG, Brazil.; Departamento de Biodiversidade Evolução e Meio Ambiente, Universidade Federal de Ouro Preto, Campus Morro do Cruzeiro s/n, Bauxita, CEP: 35400-000, Ouro Preto, MG, Brazil.

**Keywords:** interaction network, pollen transportation, territorial behavior, Trochilidae

## Abstract

Interspecific competition can strongly influence community structure and shape niche breadth and overlap. One of the main factors that determine the hummingbird community structure is competition for food. Hummingbirds functional attributes, such as beak length and body mass, influence nectar acquisition in the flowers, shaping foraging niches according to hummingbird dominance and foraging strategy. This study evaluates how the hummingbirds’ functional and behavioral attributes are related to plants assemblage in rocky outcrops’ habitats. We tested the following hypothesis: H1) Functional traits (beak length and body mass) are related to the richness and frequency of pollen grain morphotypes carried by hummingbirds; H2) Dominant and territorial hummingbirds carry a lower richness and frequency of pollen types when compared to subordinate hummingbirds, and H3) Hummingbird species carry different types of pollen grains. We conducted the study between September 2018 and March 2019 in a *Campo Rupestre* (rocky outcrops) in Southeastern Brazil. Hummingbirds were captured with a trap built based on trapdoors. We recorded their beak size and body masses, marked with commercial bird rings and ink on parts of the body, and then released. Behavioral responses to artificial feeders were collected regarding each visit’s time and duration and the outcome of aggressive interactions. The pollen adhered to the body parts was collected and identified in the laboratory. Our results showed that neither body size nor aggressive behaviors influenced pollen richness and frequency in rocky outcrops. Beak length was the most important hummingbirds’ attribute that influenced pollen richness, but not pollen frequency. Short-billed hummingbirds carried the greatest richness of pollen grains. Pollen grain richness and frequency were not related to hummingbird body mass or aggressive behavior. The hummingbird-pollen grain interaction network has shown to be generalized in the pollen grain transport. We conclude that hummingbirds’ beak length is the central morphological variable to measure pollen grain transport. It has direct implications for the pollination of different plant species.

## Introduction

Diet specialization, the use of a subset of available resources, is considered one of the major mechanisms permitting species co-occurrence through reduced niche overlap (Chesson 2000; Levine and HilleRisLambers 2009). Diet specialists often have morphological adaptations that allow them to extract resources more efficiently than species with generalist feeding morphologies (Forister et al. 2012).

Morphological attributes of hummingbirds, such as beak and tongue lengths, and flowers, such as the corolla size, are directly related to the acquisition of nectar by hummingbirds (Sick 1997, Fogden et al. 2014, Rico-Guevara et al. 2019). When feeding, hummingbirds can carry a certain amount of pollen grains glued to different parts of their bodies (Von Matter et al. 2010, Rico-Guevara et al. 2019). Flowers with a large corolla opening are efficient in depositing pollen grains in the bodies of hummingbirds (Buzato et al. 2000). Body mass is a morphological variable of great importance in hummingbirds’ interactions (Feinsinger 1976, Araya-Salas et al. 2018). Short-billed hummingbirds are generally characterized as functionally generalized pollinators, visiting many flowers with short corollas (Von Matter et al. 2010, Fogden et al. 2014, Maruyama et al. 2016). On the other hand, hummingbirds with long and curved beaks are adapted to the exploitation of flowers with long corollas (Collins & Paton 1989, Buzato et al. 2000, Fogden et al. 2014).

Body size and body mass have been related to the dominance hierarchy of hummingbirds in different assemblages. They can influence the composition of hummingbird species exploring a given food source, explaining in part the niche specialization observed in this group of birds (Wolf et al. 1976, Feinsinger & Colwell 1978, Lopez-Segoviano et al. 2018, Marquez-Luna et al. 2019). Overall, larger hummingbird species tend to dominate, excluding smaller species from high-quality energy resources (Araujo-Silva and Bessa 2010; Justino et al. 2012; Mendiola-Islas et al. 2016, Marquez-Luna et al. 2019). However, smaller species can also establish and defend foraging territories against larger contenders (Wolf et al. 1976; Antunes 2003). Hummingbirds of medium-large body masses are more efficient than… in expelling smaller species from food resources (Lanna et al. 2017, Lopez-Segoviano et al. 2018). Generally, medium-sized hummingbirds and small-medium straight beaks are territorial species, dominant in defense of the food resource (Feinsinger 1976). Thus, it is expected that medium-large body masses hummingbirds present less richness and less frequency of pollen adhered to their bodies when compared to small body masses species.

Dominance or subordination of hummingbirds can lead them to adopt two strategies for acquiring their food: 1) territorialism or, 2) trapline (Stiles 1975, Feinsinger & Colwell 1978). Hummingbirds that are subordinate and do not defend territories must forage for greater distances, visiting a more significant number of flower species, which can increase the richness and frequency of pollen grains adhered to their bodies. Since dominant hummingbirds defend and have exclusive access to more energetic food resources, they do not need to forage for greater distances, gaining more benefits than subordinate hummingbirds (Lanna et al. 2017). Dominant and more aggressive hummingbirds may have, then, less richness and less frequency of pollen grains adhered to their bodies (Lanna et al. 2017).

Thus, hummingbirds’ and plants’ morphological attributes can facilitate or hinder the relationship between them when referring to pollination (Sick 1997), which may influence the transport and transfer of pollen grain from one flower to another (Sick 1997, Von Matter et al. 2010).

The interaction between plants that provide food to birds (in the form of nectar or fruits) and birds that provide positive services to the plants (in the way of pollen transfer or seed dispersal) has attracted the attention of biologists since Darwin’s time (Bascompte and Jordano, 2007, Maruyama et al. 2016, Vizentin-Bugoni et al. 2016). Studies are now using complex interaction network metrics to describe bird-plant mutualistic interactions and are contributing to the development of the conceptual framework of ecological networks (Bascompte and Jordano, 2007; Ings et al., 2009; Heleno et al., 2014; Gu et al., 2015; Rodríguez-Flores et al. 2019, Simmons et al. 2019). A pattern is emerging from these studies: the interspecific interactions between plant-hummingbird present an uneven distribution between species (i.e., the degree of dependence of one species to another), with some hummingbirds interacting with several plant species and others interacting with few plant species (Rezende et al. 2007).

In the context presented above, in our study, we investigate whether morphological traits (beak length and body mass) and behavioral traits (dominance hierarchy) influence the richness and frequency of pollen grains transported in the hummingbirds in outcrops in southeastern Brazil. We also investigate the influence of these traits in metrics of the plants-hummingbird ecological network. We tested the following hypotheses: H1) The richness and frequency of pollen grains transported in the hummingbird body are influenced by beak length and body mass. We expected short-billed hummingbirds to carry more richness and pollen frequency because they visit more flowers than long-billed hummingbirds. H2) Dominance behavior will influence the richness and frequency of pollen grains in the hummingbird body. We expect that dominant hummingbirds will carry a lower richness and frequency of pollen grain than subordinate hummingbirds because they spend more time feeding and defending a single spot of a preferred resource. H3) The hummingbird-plant interactions will present specializations with a lower overlap in the composition of morphotypes. We expect hummingbirds’ morphological and behavioral characteristics to influence the richness and frequency of pollen carried by the birds with a lower overlap in the pollen types composition.

## Methods

### Study area

The study was conducted between September 2018 and March 2019 in a *Campo Rupestre* (rocky outcrops) site located in Ouro Preto city, Minas Gerais, Southeastern Brazil (20°22’16.62” S, 43°30’23.43” W). The altitude is 1,397m, and the climate is humid-mesothermal (humid temperate, with dry winters and hot and rainy summers; Álvares et al. 2013).

### Hummingbird capture

The study was divided into two stages: 1) capture of hummingbirds, held in September and November 2018, and 2) sampling of hummingbirds’ dominance behaviors, which were carried out in January and February 2019, in two periods (morning and afternoon), totaling 195 hours in 30 non-consecutive days. The study summed a sampling effort of 438 hours.

Hummingbirds were captured with a trap built based on trapdoors used in hunting wild birds. It consists of a crate 50-60cm wide and 50cm high. All sides of the box were closed with a fine screen (green net). On one side of 60cm, a door was kept open with a 30cm bamboo rod. A string was attached to the door’s support rod, which, when pulled, caused the door to close, trapping the hummingbirds inside.

Hummingbirds were attracted to the trap’s interior with artificial drinking fountains filled with a 25% sugar concentration solution (Lanna et al. 2017). The drinkers were offered from 6:00 am to 6:00 pm, two days before data collection. In the following three days, the hummingbirds were captured from 2:00 pm to 6:00 pm to ensure that these individuals had visited natural flowers during the morning. The sampled individuals were marked with commercial bird rings and ink on three parts of the body (chest, tail, and back). This method was preferred to minimize possible loss of pollen due to the manipulation of hummingbirds in ornithological nets traditionally used in this type of work (Borgella et al. 2001, Avalos et al. 2011). The capture of hummingbirds was authorized by the Brazilian Agency of Environment and Renewable Sources (SISBIO license number 51082-1). We measure each bird’s beak length using a digital caliper (model ZAAS 1.004®) and body mass, using a Pesola precision scale (LogNature®).

The pollen adhered to the body, forehead, throat, and beak of the individuals was removed using small portions of glycerin gelatin stained with fuchsin (Beattie 1971) and stored in Eppendorf for subsequent preparation of the slides for identification. Identification of morphotypes was performed considering the ornamentation, shape, and size of the pollen grains (Murcia & Feinsinger 1996, Fonseca et al. 2016).

### Hummingbird behavior

We documented the interactions among hummingbirds by conducting behavioral observations at a distance using an artificial drinking fountain to attract hummingbirds (following Lanna et al., 2017). The artificial drinker contained three 200 ml (Mr. Pet®) artificial flowers and was filled with a 25% sugar solution. The fountain was remained available throughout the day, and the mixture was always replaced every morning (Lanna et al. 2016, Lanna et al. 2017).

Dominance behaviors were recorded from 6 am to 10 am and from 2 pm to 4 pm on five consecutive days. Birds were observed at a distance of 8m, using a 10×50 binocular (Nikon TX Extreme), and the hummingbirds were identified according to Sigrist (2009). For each observation, we recorded the hummingbird species, the time and duration of each visit, and the outcome of aggressive interactions. The aggressive interactions were characterized by a hummingbird chasing (without contact) and/or attacking other hummingbirds (Cotton 1998, Camfield 2006). The winner was identified as the hummingbird that returned to feed or perch nearby after it had successfully defended and/or chased off another hummingbird from the feeder. For behavioral data collection, the focal sampling was used to record all agonistic behavior occurrences (Altmann 1974).

### Statistical analysis

We determined pollen grain frequency in the hummingbirds’ bodies by summarizing the abundance of pollen morphotypes in each hummingbird species divided by the number of individuals of each hummingbird species. Morphotypes with a frequency lower than 3% were removed because these types could represent possible contamination of pollen grains from a plant species deposited on the flower by any other floral visitor (Talavera et al. 2001).

Pearson’s correlation analysis was performed to verify whether the hummingbird’s beak length was correlated with its body mass. Subsequently, Generalized Linear Models were built, and, in the case of non-significant variables, the models were reduced to the minimum ideal model (Zuur et al. 2009). We created four models, two related to the hummingbird’s morphology, and the other two related to the hummingbird behavior. To analyze if the hummingbird’s morphology affected the pollen richness and the frequency of pollen morphotypes (as response variables) and the beak length and body mass as explanatory variables (the error distributions were Poisson and Gaussian, respectively). And, to analyze if the hummingbird behavior affected the pollen richness and the frequency of pollen morphotypes (the response variables), the models had the number of victories in agonistic combats and the time the species spent feeding as explanatory variables (the error distribution were negative binomial and Gaussian, respectively).

A mutualistic network was built to evaluate the relationship between the frequency of each pollen grain morphotype transported by the hummingbird. We used a quantitative matrix with hummingbirds in the columns and pollen morphotypes in the lines (Blüthgen et al. 2006). The indexes used were: complementary specialization (H2’) and modularity (Q). The H2’index is derived from Shannon entropy, which describes the diversity of interactions (distribution of the interactions’ weight in the interaction network). It is suggested to be robust for differences in the sampling effort and the size of the analyzed network (Fründ et al. 2016). Values close to 0 indicate high specialization, and values relative to 1 indicate high generalization (Newman 2006, Blüthgen et al. 2007). The Q index shows the most related species within the network, ranging from 0 to 1. Values close to 0 indicate no connection between individuals, and close to 1 indicates individuals high connected in the network (Dormann and Strauss 2014). We used the bipartite package to calculate and analyze the interaction network (Dormann et al. 2009) of the software R version 3.1.0 (R Development Core Team 2017).

## Results

Eight species of hummingbirds were recorded (Table 1)in the study. Of these, only *Phaethornis eurynome* was not observed during behavioral sampling. Pearson’s correlation analysis showed that the length of the beak (mm) and body mass (g) of hummingbirds were not correlated variables. In our study, the highest richness of pollen grain morphotypes was recorded in *Amazilia lactea* (s = 8), and the lowest richness was recorded in *Phaethornis pretrei* (s = 1) (Table 1). The highest frequency of pollen grain morphotypes was found in *Phaethornis pretrei* (n = 96.02%), and the lowest frequency in *Heliodoxa rubricauda* (n = 27.02%) (Table 1). The hummingbird species with the highest body mass was *Eupetomena macroura* (8.5g), and the lowest was *Chlorostilbon lucidus* (4.0g) (Table 1). The species with the longest beak was *Phaethornis pretrei* (33.5mm), and the shortest beak was *Amazilia lactea* (14.3mm) (Table 1).

**Table 1:**
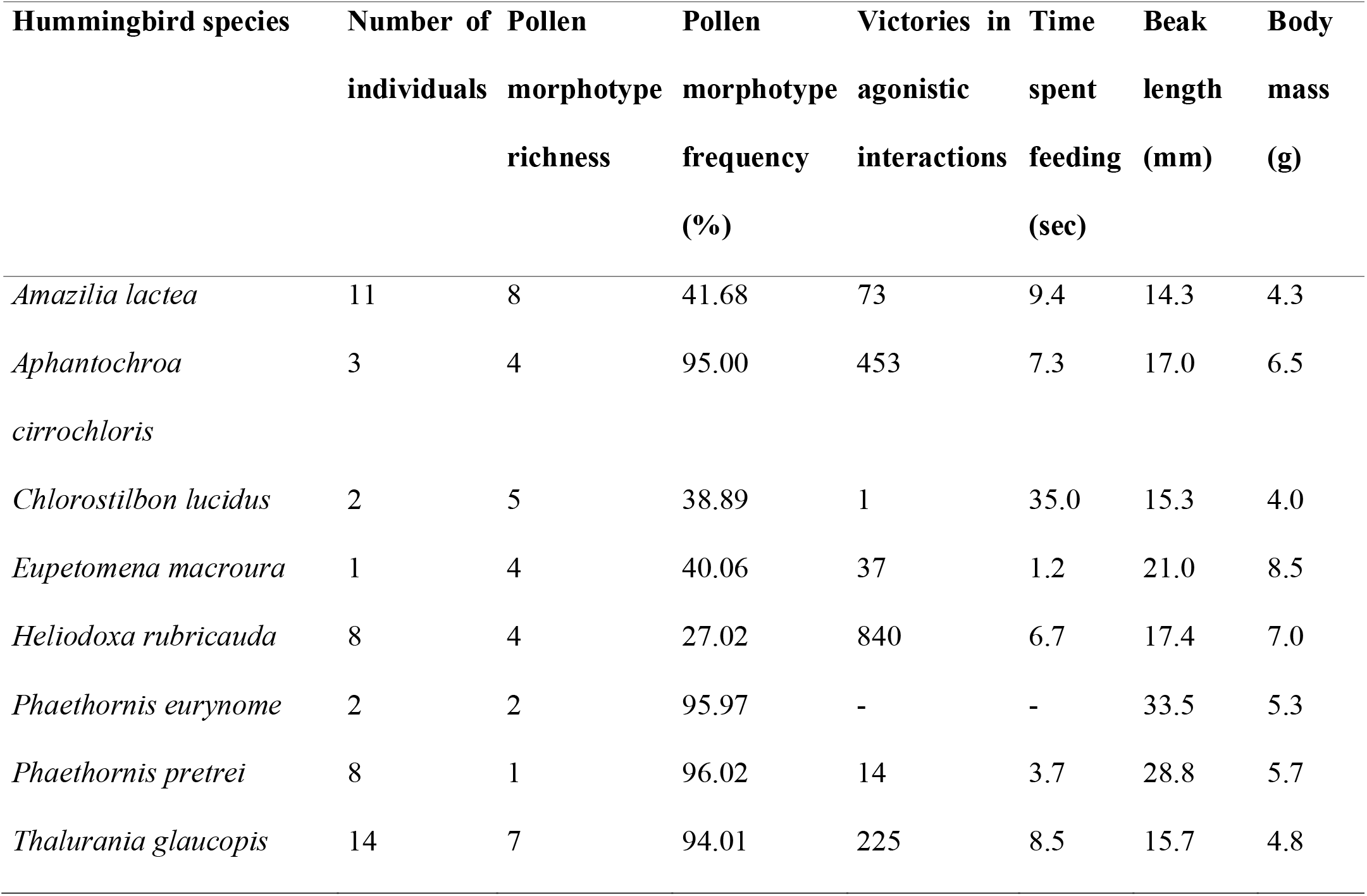
Physical attributes of the recorded hummingbirds, average time spent feeding, and the number of victories in the agonistic interactions while serving in the drinking fountain. Richness and frequency of pollen grains morphotypes transported by each hummingbird species.

Forty-seven morphotypes of pollen-grains adhered to the hummingbirds’ bodies were identified. With the removal of the morphotypes that had a frequency below 3%, the final richness was 15 pollen grains morphotypes. In this study, the species that obtained the highest number of victories in interspecific and intraspecific agonistic combats was *Heliodoxa rubricauda* (n = 840), and the least amount was *Chlorostilbon lucidus* (n = 1) (Table 1). The species that spent the longest time feeding was *Chlorostilbon lucidus* (35 sec), and the shortest time was *Eupetomena macroura* (1.3 sec) (Table 1).

The richness of pollen grain morphotypes was inversely related to the length of the hummingbird beak (F_1, 6.3218_ = 25.755, p <0.004) (Figure 1; Table 2) and did not vary with hummingbirds’ body mass (F_1, 0.008_ = 0.0287, p <0.873). The frequency of pollen grain morphotypes did not vary significantly depending on the length of the beak (F_1, 828.07_ = 0.8126, p <0.409) and on the body mass of hummingbirds (F_1, 521.35_ = 0.4560, p <0.536) (Table 2).

**Figure 1:**
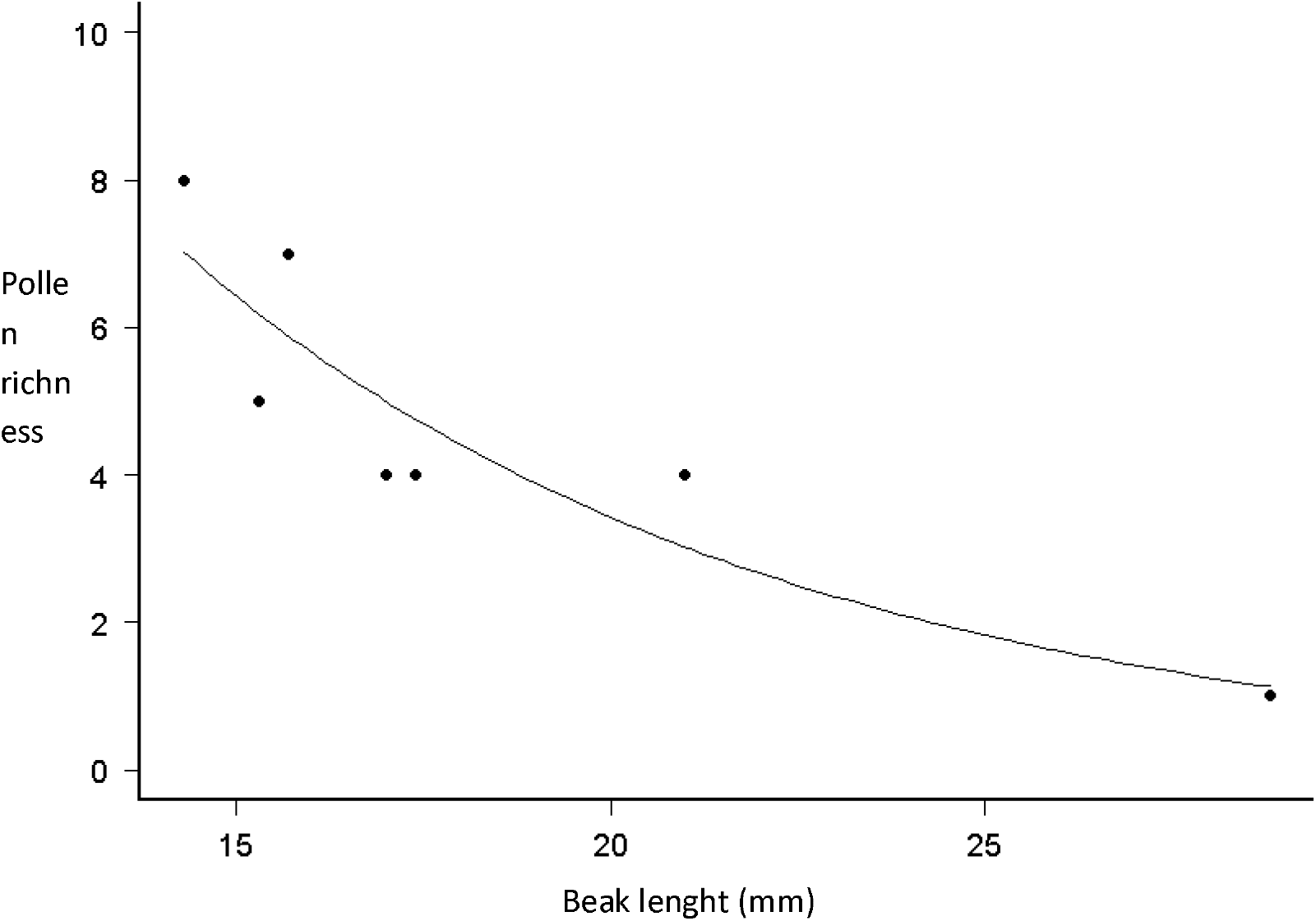
Relationship between richness of pollen grains morphotypes carried by hummingbirds and the length of the beak.

**Table 2:**
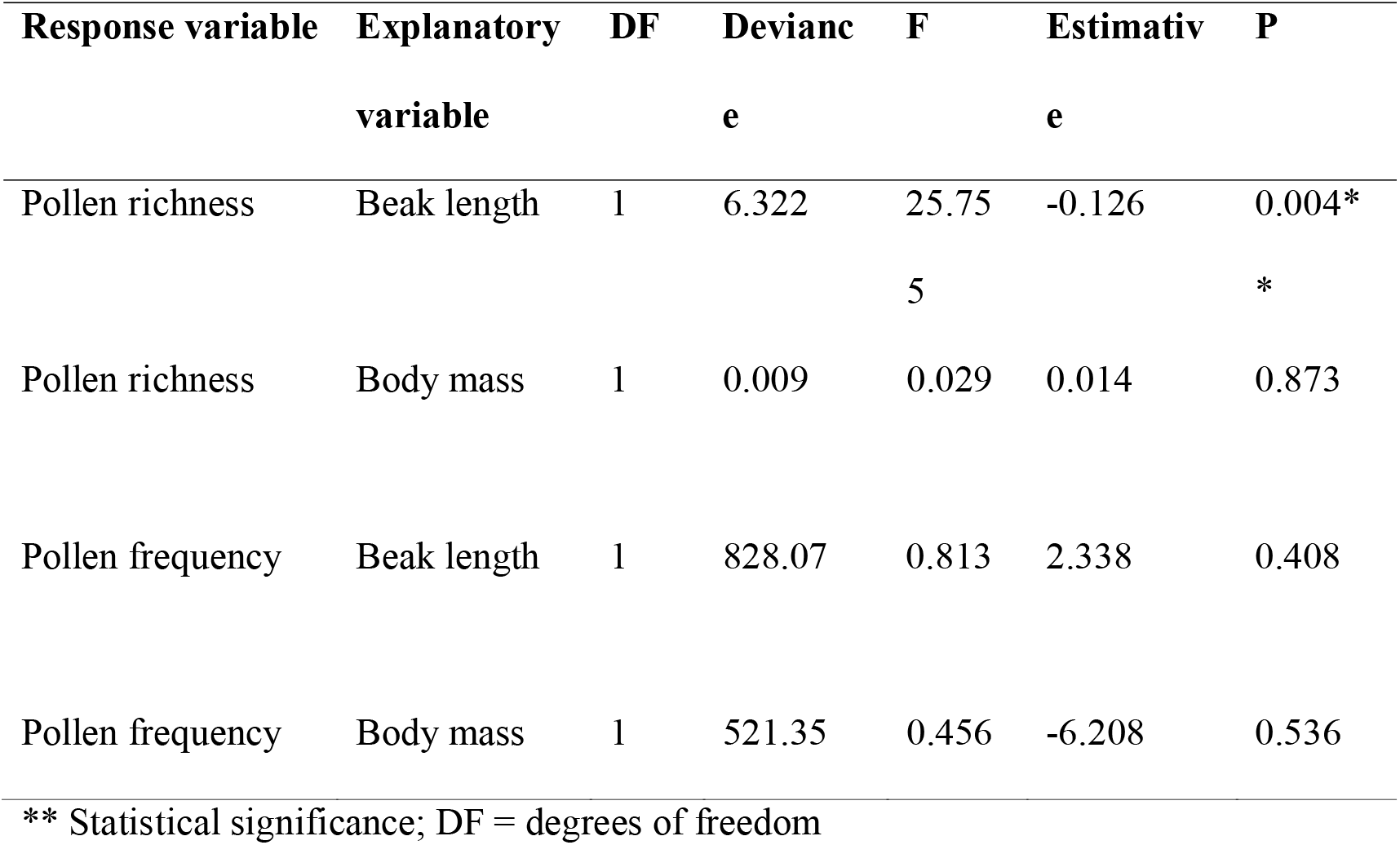
Results of the GLM models for richness and abundance of pollen grains morphotypes collected in hummingbirds according to the functional attributes body mass plus beak length.

Dominance behaviors in the artificial feeder did not vary with the richness and frequency of pollen grain morphotypes found in hummingbirds (Table 3). Hummingbirds’ body mass also did not change concerning the number of victories in agonistic interactions on artificial feeders (Table 3).

**Table 3:** Richness and frequency of pollen grain morphotypes as a function of the total time spent in feeding and the number of victories in agonistic interactions.

| **Response**<br>**variable** | **Explanatory**<br>**variable** | **DF** | **Deviance** | **F** | **Estimative** | **P** |
| --- | --- | --- | --- | --- | --- | --- |
| Pollen frequency | Time spent feeding | 1 | 446.17 | 0.407 | 2.410 | 0.551 |
| Pollen frequency | Victories | 1 | 131.83 | 0.106 | -0.033 | 0.760 |
| Pollen richness | Time spent feeding | 1 | 8.696 | 1.913 | 0.336 | 0.225 |
| Pollen richness | Victories | 1 | 0.105 | 0.021 | -0.002 | 0.890 |

DF = degrees of freedom

The interaction network between hummingbirds and pollen grain morphotypes showed median values of modularity (Q= 0.533) (Figure 2) and the complementary specialization index H2’(0.675) (Figure 3), indicating medium connectivity between hummingbirds and plants and a tendency for generalization of these relations.

**Figure 2:**
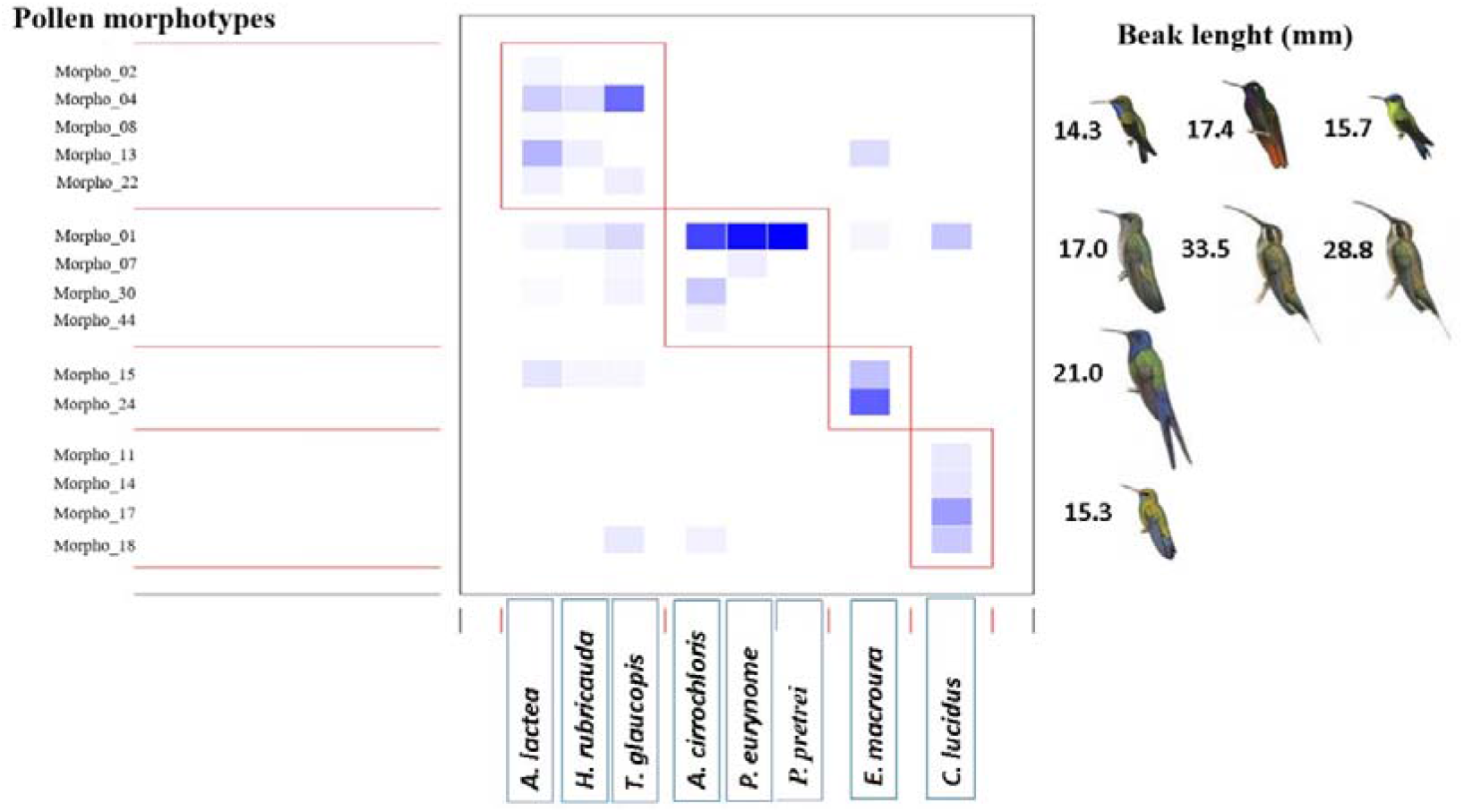
Interaction network (modules) formed by the hummingbird species and the pollen grains morphotypes concerning their beak lengths. The most vibrant colors represent the highest frequency of pollen carried by them.

**Figure 3:**
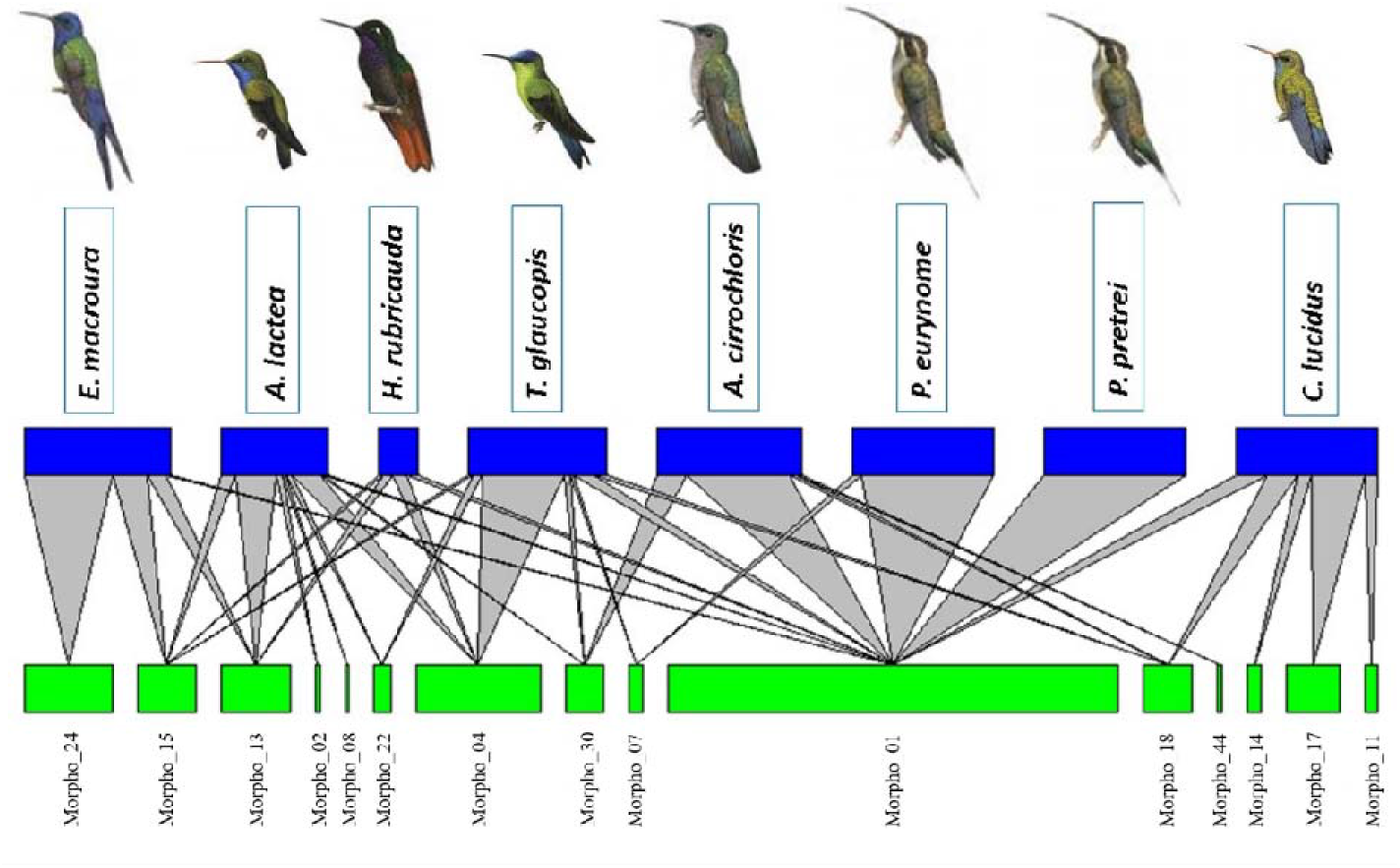
Interaction network representing the overlapping trophic niche of hummingbirds based on the frequency of pollen grain transported by them.

## Discussion

Our results showed that neither body size nor aggressive behaviors influenced pollen richness and frequency in rocky outcrops. Beak length was the most crucial hummingbirds’ attribute that influenced pollen richness, but not pollen frequency. We expected short-billed hummingbirds to carry a higher richness of pollen morphotypes with higher frequency because they need to visit more flowers than long-billed hummingbirds. We found that the length of the beak of hummingbird species proved to be the most important morphological trait related to the richness of plants visited (measured by the pollen grains richness) and less important to explain the frequency of pollen grains transported. This greater richness of pollen grains is probably the result of an adjustment of the hummingbirds’ beak with the flowers they feed on (Sick 1997, Fogden et al. 2014, Rico-Guevara et al. 2019). Due to that adjustment, the interaction network between hummingbirds and pollen grains was moderately specialized, reflecting the importance of hummingbirds in different plants’ pollination process.

The short-billed hummingbird species in our study (*A. lactea, T. glaucopis,* and *C. lucidus)* carried a higher pollen grain richness. These species are considered functional generalists because they can visit a great diversity of native and exotic plants (Maruyama et al. 2016), mainly plants with short corollas (Maruyama et al. 2016).). At the other extreme, the longest, curved-billed hummingbird species (*P. eurynome* and *P. petrei)* carried a lower richness of pollen grains when compared to the intermediatebilled species (*A. cirrochloris, H. rubricauda,* and *E. macroura)* and to the short-billed (*A. lactea, T. glaucopis,* and *C. lucidus).* These two *Phaethornis* hummingbirds are considered specialized in feeding nectar from long tube bromeliad flowers found in the Atlantic forest, restricting access to nectar for short-billed species hummingbirds (Buzato et al. 2000, Maruyama et al. 2016, Sonne et al. 2019). The morphological match and phenological overlap are important factors predicting plant-hummingbird interactions, showing the role of these characteristics in organizing plant-hummingbird communities (Brown & Bowers 1985, Vizentin-Bugoni et al. 2014). Thus, the lowest richness of pollen grains found in *Phaethornis* species may reflect the lack of long, tubular flowers in the study area, remembering that our study was conducted in the *Campo Rupestre* area where more plant species have open and narrow corollas (Giulietti et al. 1997).

On the other hand, the frequency of pollen grains was not significantly related to the beak length. Our results indicate that the potential pollinators’ effectiveness may be mediated more closely by the abundance of pollinators (hummingbirds) than the trait-matching compatibility. The potential pollinators are the pollinator’s total contribution to the plant’s fitness, measure, for example, by the number of pollen grains deposited (Schupp et al. 2017, Missagia and Alves 2018). Although hummingbirds’ beak was not a good predictor to explain a higher frequency of pollen grains in our study, the result points to a tendency for long, curved-billed species to carry a higher frequency of pollen grains than short-billed species. However, it is crucial to consider the environment where the present study was carried out. A rocky outcrop is a less complex and more unpredictable habitat; although deemed a megadiverse, it presents less flower diversity than forest habitats (Silveira et al. 2016). Maybe, more complex, flower-diverse habitats, like forests and urban environments, could increase the frequency of pollen carried by short-billed hummingbirds, due mainly to differences in plant (flower) composition (Maglianesi et al. 2014).

The richness and frequency of pollen grain morphotypes were not influenced by the hummingbirds’ body masses, as expected. Lighter hummingbirds, such as the Phaethornithinae *(P. eurynome* and *P. pretreí)* and the Trochilinae *(C. lucidus, A. lactea,* and *T. glaucopis)* were expected to carry more morphotypes and more pollen grains. Species that are subordinate avoid combats with more dominant species. Therefore, they need to visit more plant species, which would increase the chances of having more pollen of different plant species in their bodies. Thus, hummingbirds’ body mass and dominance behavior were not variables that explained the differences in the number of pollen grains transported. In contrast, some authors cite that the behavior and the body mass are the most important variables for structuring the hummingbird assembly in a resource spot (Sick 1997, Forgden et al. 2014, Lanna et al. 2017). More aggressive and dominant species can share the territory with more peaceful and less hostile species (Stiles 2008, Lanna et al. 2017, Lopez-Segoviano et al. 2018). Marques-Luna et al. (2019) found that the dominant species of hummingbirds in the assemblage *(L. clemenciae* and *C. thalassinus)* weigh more than 6 g, representing the largest hummingbird species in the community. The higher rank within the dominance hierarchy was associated with large body size species, which coincides with that reported by different authors (Dearborn 1998; Justino et al. 2012). However, the bigger species are not always the most aggressive and dominant, observed by Martin and Ghalambor (2014) and Márquez-Luna et al. (2019), where smaller species of hummingbirds dominated bigger ones in some of the interactions.

The interaction network between hummingbirds and pollen grain morphotypes tended to be generalized, with an overlap of pollen grain morphotypes carried by each hummingbird species. Most of the hummingbirds recorded in the present study belonged to the Emerald clade *(Chlorostilbon, Eupetomena, Amazilia, Thalurania,* and *Aphantochroa),* formed by evolutionarily recent and more generalist hummingbirds. The *Phaetornis* genus belongs to the Hermit clade, formed by evolutionarily older and more specialized hummingbirds (Rodrigues-Flores et al. 2019). Because of the hummingbird species involved, we expected and observed in this study a heterogeneous connectivity distribution, with a tendency to generalize the species interaction in the network, because most hummingbirds visited more plant species and few hummingbirds visited few of the plants.

Hummingbirds present a preference in selecting their food resource (Stiles 1975, Feinsinger et al. 1979, Maruyama et al. 2014, Lanna et al. 2017). This specialization can be related to the flower morphology (Maglianesi et al. 2014, Fonseca et al. 2016) and to hummingbird morphology (beak length) (Maglianesi et al. 2014). However, it was shown that hummingbirds relaxed their specialized relationships and became less specific in their interactions in non-forested areas (Morrison and Mendenhall, 2020). Our research was conducted in a *Campo Rupestre* area, where trees are rare, reinforcing our result of a lower specialization in our interaction network. Thus, most hummingbirds explored many flowers, ornithophilous or not, while the hermits remained to explore few plant species.

We conclude that the hummingbirds’ beak’s length is the most important variable to explain the richness, specialization, and segregation in the transport of pollen grains by hummingbirds in rocky outcrops. As suggested by Maglianesi et al. (2014), hummingbird species’ morphological traits influence ecological specialization patterns. The beak’s morphology seems to be more important than body mass in determining the niche partition within the community. In this way, the beak of hummingbirds is of great importance for the pollination process of many species of plants.

## Acknowledgments

We are grateful to CAPES for the scholarship granted to RMC and MVB and YA’s financial support. We are thankful to UFOP’s Biodiversity laboratory for providing all the logistics for data collection. We also thank Dr. Fernanda Costa for her assistance with the interaction network analysis.

